# Disrupting the ArcA regulatory network increases tetracycline susceptibility of Tet^R^ *Escherichia coli*

**DOI:** 10.1101/2020.08.31.275693

**Authors:** Mario L. Arrieta-Ortiz, Min Pan, Amardeep Kaur, Vivek Srinivas, Ananya Dash, Selva Rupa Christinal Immanuel, Nitin S. Baliga

**Author notes:** corresponding author, Nitin S. Baliga, 401 Terry Ave N, Seattle, WA 98109, (NSB).

## Abstract

There is an urgent need for strategies to discover secondary drugs to prevent or disrupt antimicrobial resistance (AMR), which is causing >700,000 deaths annually. Here, we demonstrate that tetracycline resistant (Tet^R^) *Escherichia coli* undergoes global transcriptional and metabolic remodeling, including down-regulation of tricarboxylic acid cycle and disruption of redox homeostasis, to support consumption of the proton motive force for tetracycline efflux. Targeted knockout of ArcA, identified by network analysis as a master regulator among 25 transcription factors of this new compensatory physiological state, significantly increased the susceptibility of Tet^R^ *E. coli* to tetracycline treatment. A drug, sertraline, which generated a similar metabolome profile as the *arcA* knockout strain also synergistically re-sensitized Tet^R^ *E. coli* to tetracycline. The potentiating effect of sertraline was eliminated upon knocking out *arcA*, demonstrating that the mechanism of synergy was through action of sertraline on the tetracycline-induced ArcA network in the Tet^R^ strain. Our findings demonstrate that targeting mechanistic drivers of compensatory physiological states could be a generalizable strategy to re-sensitize AMR pathogens to lost antibiotics.

## INTRODUCTION

Antimicrobial resistance (AMR) is the ability of a bacterium to withstand growth inhibition and killing by high doses of an antibiotic^1,2^. The problem of AMR has emerged from the over-prescription and overuse of antibiotics^3–5^, accumulation of antibiotics in the natural environment^6^, antibiotic-induced increased mutation rates^7,8^, horizontal transfer of resistance-conferring genes^3^, and poor infection control strategies^9^. As a result, infections by pathogenic AMR strains is rapidly growing and projected to cause ~10 million deaths by 2050^4^. While health policies to regulate antibiotic use^10^ and programs to ensure patient compliance with completing prescribed antibiotic regimens are effective^11^, these efforts are expensive, laborious, and face implementation challenges across the world^12^. Similarly, efforts to find new potent antibiotics are effective^13^ but also expensive and being outpaced by the rate at which new resistant strains are emerging^14^.

A solution to tackling AMR might be in the observation that gain of resistance to an antibiotic is typically associated with loss of fitness^3^, which is restored through compensatory mutations^3^ and changes in regulation and metabolism^15,16^. For example, *Pseudomonas aeruginosa* upregulates anaerobic nitrate respiration to quench intracellular protons and compensate for loss of fitness due to efflux-mediated resistance^17^. Similarly, *Mycobacterium smegmatis* transcriptionally up regulates the rRNA methylase TlyA to restore fitness upon gaining resistance to capreomycin^18^. Molecules that target new vulnerabilities within these compensatory mechanisms required to support the gain of antibiotic resistance could enable the recovery of “lost” antibiotics and prolong the lifespan of new antibiotics^19,20^. The ability of metabolite supplementation to re-sensitize resistant pathogens to diverse antibiotics, including aminoglycosides^21^, chloramphenicol and streptomycin^22^ lends credibility to this idea. However, to implement such a strategy at scale we need to develop methodology to discover the mechanistic driver of fitness-restoring compensatory changes in AMR strains, confirm with targeted genetic perturbations that it indeed represents a new vulnerability, and use a rational approach to find molecules that could disrupt the compensatory mechanism^23^.

Here, we have developed a systems approach to discover molecules that can synergistically resensitize tetracycline-resistant *E. coli* (hereafter Tet^R^) to tetracycline. Discovered in 1947, tetracycline is a protein synthesis inhibitor that acts by binding to the 30S ribosomal subunit^24,25^. Tetracycline was rapidly adopted in the clinic due to its efficacy against a broad spectrum of pathogens^24,25^, and continues to be used widely in animal farming^26^. Resistance to tetracyclines emerged a few years later (1953) and progressively reduced their effectiveness^24,25^. The primary mechanisms of tetracycline resistance are: i) through active extrusion by efflux pumps; ii) gain of mutations that disrupt interaction with the target; and iii) enzymatic inactivation, e.g., by TetX^24,25^. Previous attempts to counteract tetracycline resistance in *E. coli* have focused on potential efflux pump inhibitors^27,28^.

We have discovered that when *E. coli* gains resistance to tetracycline through AcrAB-mediated efflux, a global shift in metabolism to a fermentative state is required to restore fitness of the resistant strain in the presence of tetracycline. The regulatory network that mechanistically drives this global metabolic reprogramming in the Tet^R^ strain is comprised of at least 25 TFs that directly regulate 279 genes. Interestingly, 209 of the 279 genes are differentially regulated by 15 TFs only in the presence of tetracycline, suggesting changes in activity of these regulators by downstream consequences of the increased activity of the AcrAB efflux pump. Knocking out *arcA*, a master regulator of this network, significantly reduced the fitness of the Tet^R^ strain. We discovered that the drug sertraline, which generated a similar metabolome profile as the ArcA knockout, synergistically potentiated the bacteriostatic effect of tetracycline on the Tet^R^ strain, but not the wild type strain, on which the effect was antagonistic. Finally, we show that deleting ArcA abolished the potentiating effect of sertraline, demonstrating that the mechanism of its synergy with tetracycline was through its action on the tetracycline-induced ArcA-regulated network. We discuss these results from the perspective of formulating multidrug regimen using a network-based approach to recover lost antibiotics and prolong the utility of new antibiotics.

## RESULTS

### A novel physiological state underlies tetracycline resistance in E. coli

To characterize the direct and compensatory physiological changes triggered by the acquisition of antibiotic resistance, we reanalyzed transcriptomes of the tetracycline-susceptible wild type (MG1655, here onwards “WT”) and a laboratory-evolved Tet^R^ strain of *E. coli* with and without tetracycline treatment^29,30^. In the absence of tetracycline, the Tet^R^ strain differentially expressed 197 genes (DEGs) relative to the WT strain, including 65 metabolic genes, seven transcription factors (TFs) and four efflux pump (EP)-related genes (including the *acrAB* operon) (**Fig. 1A**). Function enrichment of the dysregulated gene set revealed that 14 pathways were significantly perturbed in the Tet^R^ strain (**Fig 1B, Table S1**). Of note was the differential regulation of 33 fermentation-related genes (randomized permutation test p-value < 0.01), including the *frd* operon, *adhE, fumC*, and *ldhA* (p-value < 0.05) (**Fig. S1**). Notably, upregulation of the *acrAB* operon and *acrZ* is consistent with known mechanisms of resistance to tetracycline and other antibiotics (**Fig. S2A**)^28,31,32^. Disruption of the AcrAB efflux pump exacerbated the sensitivity of *E. coli* to tetracycline^33^ (**Fig. S2B**). Upon re-sequencing and comparing the genomes of the Tet^R^ and WT strains, we discovered that gain of tetracycline resistance could be due to mutations in *acrB* and *acrR*, a transcriptional repressor of the *acrAB* operon, which is consistent with upregulation of the efflux pump^34^. Notably mutations in these genes may also confer resistance to chloramphenicol^35,36^.

**Fig. 1.**
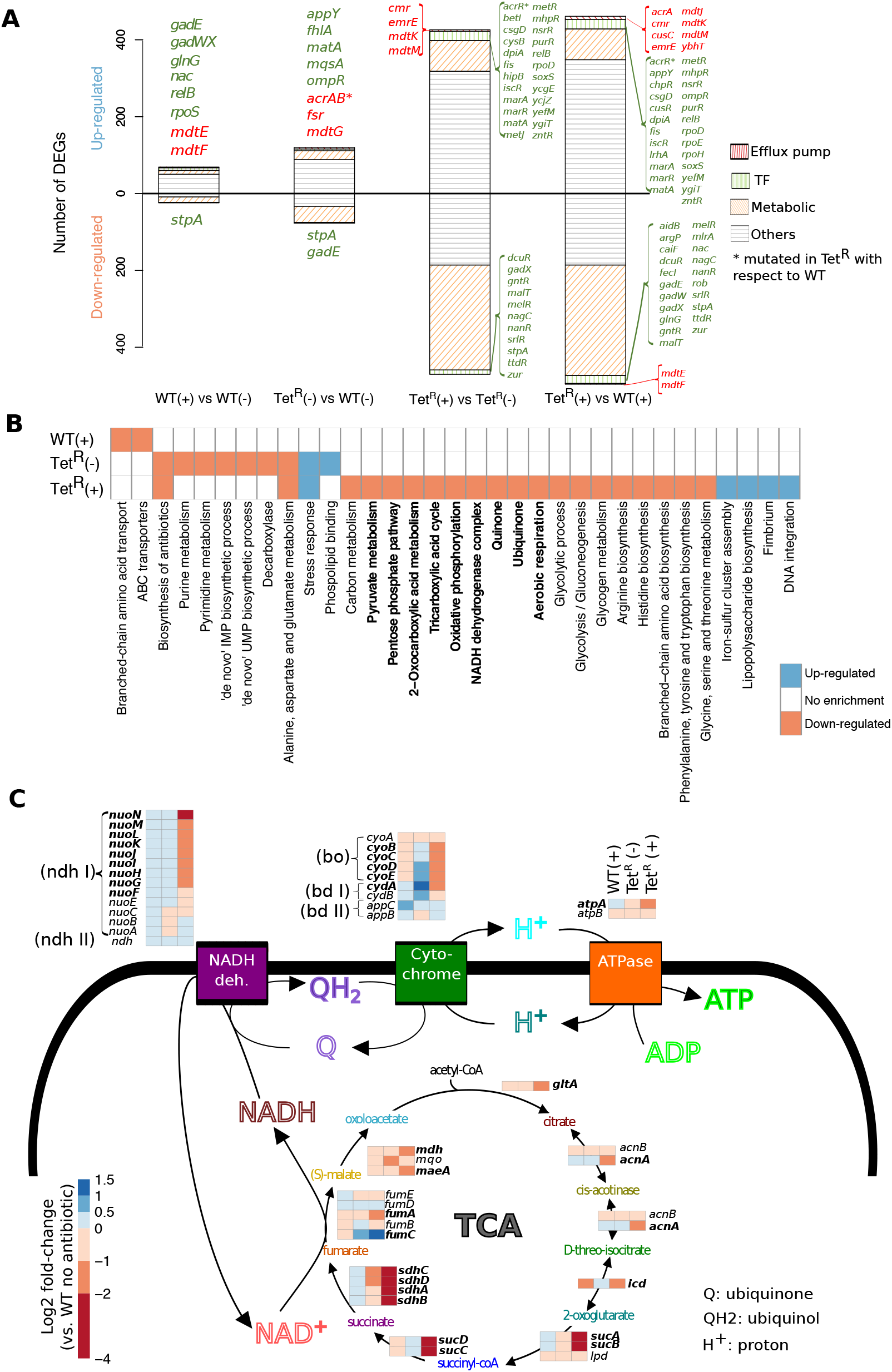
Transcriptional and metabolic remodeling accompanying gain of tetracycline resistance in *E. coli*. (A) Transcriptomes comparison for the Tet^R^ and parental WT (MG1655) strains in the presence of and absence of tetracycline (indicated as ‘(+)’ and ‘(-)’ next to the strain’s name, respectively). Analyzed data was from Händel et al^30^. Differentially expressed genes (adjusted p-value < 0.05 & absolute log2 fold-change > 1) were classified as efflux pump-related (compiled from the EcoCyc database and available literature), transcription factors (TFs; based on the transcriptional regulatory network compiled from the RegulonDB database) or metabolism-related (based on the iJO1366 metabolic model of *E. coli*)^37,73,74^. Unassigned genes were grouped in the *Others* category. Up- and down-regulated TFs and metabolic genes are listed in green and red font, respectively. (B) Heatmap with functional enrichment information of the set of genes significantly up- and down-regulated in the WT strain in the presence of tetracycline, and the Tet^R^ strain without and with tetracycline with respect to the WT strain in antibiotic free condition. Due to space constraints and functional terms redundancy, only a subset of functional terms are displayed. Functional terms shown in bold are associated with carbon metabolism and oxidative phosphorylation. (C) Fold-change profiles (with respect to the WT strain in antibiotic free condition) of genes related to the tricarboxylic acid (TCA) cycle, the electron transport chain and ATP synthase.

**Fig. 2.**
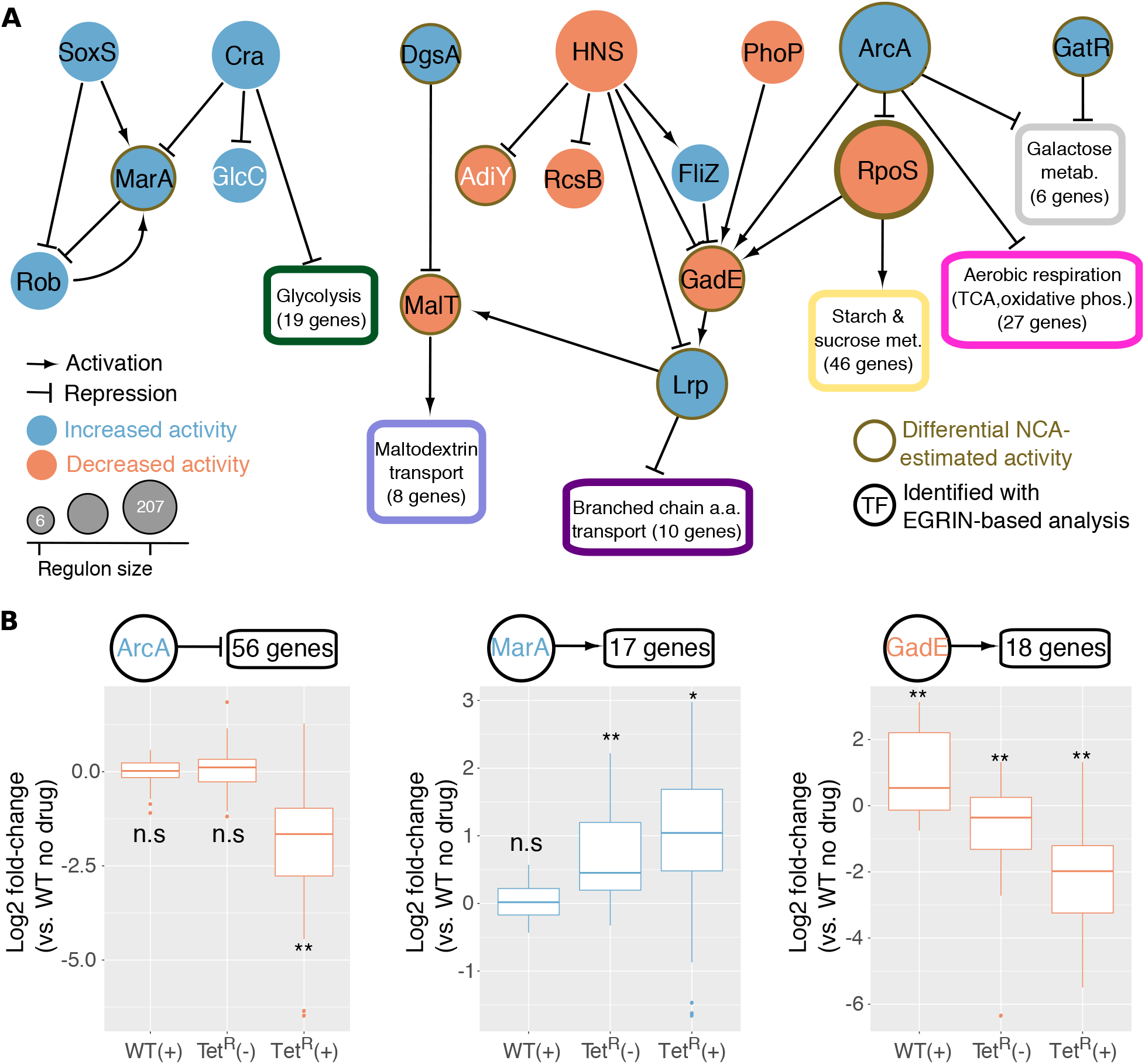
Regulatory circuits differentially active in tetracycline resistant *E. coli*. (A) Subnetwork of 17 TFs (displayed as circles) implicated in driving transcriptional and metabolic reprogramming in the Tet^R^ strain. Eight differentially active TFs that were also implicated as important in the Tet^R^ strain are not included due to lack of interactions with the 17 TFs. Function enrichment within sub-sets of 6 more genes regulated by the same TF(s) are shown within boxes. The numbers in parentheses indicate the number of genes used to perform the functional enrichment analyses with DAVID^70^. GatR was included due to the overlap between the ArcA and GatR regulons. TFs implicated by network component analysis (NCA) in regulatory and metabolic reprogramming of the Tet^R^ strain are indicated in nodes with a brown-colored border. Black font indicates TFs that were implicated based on significant overlap of their differentially regulated targets (in response to tetracycline treatment) within co-regulated gene modules in EGRIN^41^. (B) Fold-change of the ArcA, MarA and GadE regulons support their predicted increased (for ArcA and MarA) and decreased (for GadE) activity in mediating transcriptional reprogramming of the Tet^R^ strain at baseline and during adaptive response to tetracycline treatment. Boxplots indicate differentially expressed regulon members in the basal and adaptive states (total number of regulated genes by each TF is indicated above each corresponding boxplot). Absence and presence of tetracycline treatment is indicated with ‘(-)’ and ‘(+)’, respectively. Statistical significance of the observed mean fold changes was evaluated by determining the null distribution of mean fold change in 10,000 random samplings of gene sets of similar size. p-values are indicated with * (<0.05) and ** (<0.001). N.S. indicates no significant difference.

The Tet^R^ strain differentially expressed nearly ten times as many genes as the WT (896 DEGs, adjusted p-value < 0.05 and absolute log2 fold change >1) in response to tetracycline at 0.25x minimum inhibitory concentration (MIC; 16 ug/ml) (**Fig. 1A**)^30^. This differential regulation represented reprogramming of 75 pathways, including the tricarboxylic acid (TCA) cycle (15 of 21 genes; hypergeometric test p-value = 1e-06), the electron transport chain (ETC, 14 of 23 genes; hypergeometric test p-value = 3.9e-05) and ATP synthase (four of nine genes; hypergeometric test p-value = 0.1) (**Fig 1B and C, Table S1**). While the expression of the *acrAB* efflux pump did not increase further (**Fig. S2A**), at least four additional efflux pump genes were significantly upregulated with tetracycline treatment. This suggested that the large scale transcriptional remodeling, which was potentially mediated by 35 differentially expressed TFs, might constitute a compensatory physiologic state that is triggered by increased efflux pump activity in the presence of tetracycline to ameliorate the loss of fitness associated with the resistance phenotype of the Tet^R^ strain (**Fig. 1 and Fig. S3**)^3^. Specifically, repression of aerobic oxidative phosphorylation and induction of fermentation pathways suggested that a shift towards an anoxic physiologic state might be necessary to support the tetracycline resistance phenotype.

### A transcriptional program governed by 25 TFs underlies the physiological state required for tetracycline resistance

We analyzed gene expression changes induced by gain of tetracycline resistance in the context of the transcriptional regulatory network to discover mechanisms responsible for regulatory and metabolic reprogramming of the Tet^R^ strain. We compiled a signed transcriptional regulatory network of *E. coli* based on curated activator or repressor attributes to every TF-target gene interaction in RegulonDB^37^. We then used the NetSurgeon algorithm^38^ to identify within this transcriptional regulatory network the subset of TFs whose simulated overexpression and knockout explained the overall gene expression changes induced by gain of mutations (e.g., the *acrR* mutation) and treatment with tetracycline in the Tet^R^ strain. We hypothesized that the TF networks that were differentially activated in the Tet^R^ strain upon tetracycline treatment were likely to be the mechanistic drivers of the compensatory physiologic state to support the resistance phenotype. Altogether, the NetSurgeon analysis implicated 25 TFs in mechanistically driving the differential regulation of 279 genes, of which 209 genes were regulated by a subset of 15 TFs in response to tetracycline treatment (**Table 1**). Seventeen of the 25 TFs regulated each other in two TF-TF subnetworks suggesting coordination across their regulatory networks (**Fig. 2A**). While one subnetwork included TFs that were previously linked to AMR (MarA, SoxS and Rob^39^), the other subnetwork was made up of TFs (Lrp, MalT GatR, ArcA) of metabolic pathways.

**Table 1.** TFs implicated in reprogramming transcriptional response of the Tet^R^ strain.

| Transcription factor | Locus tag | Differential activity<sup>a</sup> | Response<sup>b</sup> | Regulon size<sup>c</sup> | Targets-basal<sup>d</sup> | Targets-adaptive<sup>e</sup> |
| --- | --- | --- | --- | --- | --- | --- |
| RpoS | b2741 | Decreased | Adaptive | 207 | 20 | 69 |
| ArcA | b4401 | Increased | Adaptive | 167 | 9 | 66 |
| HNS | b1237 | Decreased | Basal | 146 | 18 | 33 |
| Cra (FruR) | b0080 | Increased | Adaptive | 76 | 2 | 43 |
| Lrp | b0889 | Increased | Both | 64 | 9 | 32 |
| PhoP | b1130 | Decreased | Basal | 49 | 9 | 15 |
| GadE | b3512 | Decreased | Both | 36 | 9 | 14 |
| SoxS | b4062 | Increased | Basal | 33 | 7 | 13 |
| MarA | b1531 | Increased | Basal | 33 | 11 | 11 |
| RcsB | b2217 | Decreased | Basal | 33 | 7 | 10 |
| PurR | b1658 | Increased | Basal | 31 | 10 | 10 |
| Rob | b4396 | Increased | Basal | 22 | 8 | 7 |
| FliZ | b1921 | Increased | Basal | 20 | 6 | 6 |
| CytR | b3934 | Increased | Adaptive | 13 | 0 | 8 |
| OmpR | b3405 | Increased | Both | 13 | 4 | 6 |
| TorR | b0995 | Decreased | Basal | 12 | 3 | 4 |
| MalT | b3418 | Decreased | Adaptive | 10 | 0 | 8 |
| DgsA | b1594 | Increased | Adaptive | 10 | 0 | 7 |
| NrdR | b0413 | Decreased | Adaptive | 9 | 1 | 4 |
| AdiY | b4116 | Decreased | Both | 8 | 3 | 5 |
| YbjK | b0846 | Increased | Adaptive | 8 | 1 | 5 |
| PspF | b1303 | Increased | Both | 7 | 4 | 5 |
| GatR | b4498 | Increased | Adaptive | 6 | 0 | 6 |
| GlcC | b2980 | Increased | Basal | 6 | 3 | 3 |
| BirA | b3973 | Decreased | Adaptive | 5 | 0 | 5 |

<sup>a</sup>“Increased” or “Decreased” indicates NetSurgeon-inferred change in the activity of each TF that explains the observed transcriptional response in a given treatment.
<sup>b</sup> “Basal” indicates the role of a TF in reprogramming transcriptional response of the Tet<sup>R</sup> strain in the absence of tetracycline, whereas “Adaptive” indicates that the TF mediates transcriptional response of the Tet<sup>R</sup> strain to tetracycline.
<sup>c</sup>Total number of genes directly regulated by each TF.
<sup>d</sup>Number of DEGs regulated by a TF in the absence of tetracycline.
<sup>e</sup>Number of DEGs regulated by a TF in the presence of tetracycline.

In a second independent approach, we used Network Component Analysis to estimate the differential regulatory activity of TFs in Tet^R^ and WT with and without tetracycline treatment^40^ (*see methods).* The estimated regulatory activities of TFs were consistent with NetSurgeon predictions for nine of the seventeen TFs (indicated with brown node border in **Fig. 2A**). Finally, in a third approach we discovered that 287 DEGs were statistically enriched across 34 gene modules regulated by fifteen (of the 17 TFs) within the previously developed *E*nvironment and *G*ene *R*egulatory *I*nfluence *N*etwork (EGRIN) model for *E. coli^41^.* In summary, of the 25 TFs identified by NetSurgeon, eight were also identified by the two orthogonal approaches. Altogether, the 15 TFs implicated in the response of Tet^R^ to tetracycline collectively regulated 23.3% (209 genes, hypergeometric test p-value < 1e-26) of all DEGs, including 6 additional TFs, explaining how the response might have propagated to other genes in the genome. Notably, the predicted increased and decreased activity of TFs driving the transcriptional response of Tet^R^ strain to tetracycline, were consistent with the changes in expression profiles of their corresponding regulons across strains and treatments (**Fig. 2B**).

ArcA, a global transcriptional regulator that is typically induced under microaerobic conditions^42^ was implicated by all three approaches as a mechanistic driver of the tetracycline response in the Tet^R^ strain. ArcA is a master regulator of one of the two TF-TF subnetworks, directly regulating 66 DEGs, and influencing regulation by at least 4 downstream TFs. There was significant overlap between DEGs in the Tet^R^ response to tetracycline and the set of DEGs in an *arcA* deletion strain in anaerobic conditions^43^ (hypergeometric test p-value < 1e-11). Importantly, ArcA is a known repressor of most genes of the TCA cycle and ETC^44^ (**Fig. 2A**), and both processes were significantly downregulated in the Tet^R^ strain in the presence of tetracycline, which could have potentially perturbed NADH/NAD ratio and disrupted energy production via aerobic respiration. ArcA directly coordinates repression of the TCA cycle with activation of overflow metabolism^45^, which is a fermentation mechanism to generate energy, albeit at lower efficiency, to cope with changes in demand for protein, energy, and biomass production under changing growth conditions^46,47^. Interestingly, fermentation genes were expressed at a higher level in the Tet^R^ strain even without tetracycline treatment, although the expression of TCA genes was downregulated only in the presence of tetracycline (**Fig 1C** and **Fig. S1**). Based on these observations we hypothesized that fitness loss associated with gain of efflux-mediated resistance to tetracycline in the Tet^R^ strain is compensated by an ArcA-mediated shift to energy production by overflow metabolism^45^.

### ArcA activity ameliorates the fitness cost of tetracycline resistance

We investigated the importance of ArcA activity for efflux-mediated tetracycline resistance by constructing *arcA* knockout strains (*ΔarcA*) in the WT and Tet^R^ strain backgrounds. We performed ScanLag analysis^48^ to compare colony growth characteristics of both strains on solid LB medium. Deletion of *arcA* significantly extended the time of appearance of colonies by 38% (t-test p-value < 1e-7) in the Tet^R^ background but had little to no effect in the WT background (**Fig 3A**). This result was reproduced in liquid cultures wherein fitness was measured as the *A*rea *U*nder the growth *C*urve^49–51^, demonstrating that the tetracycline resistance mutations had increased the importance of ArcA relative to its role in the WT background. Both lag phase and loss of fitness of the Tet^R^ Δ*arcA* strain increased with tetracycline treatment at high doses (16 ug/ml-20 ug/ml) and the effects were completely reversed by complementation with an episomal copy of *arcA* (**Fig. 3B and C**).

**Fig. 3.**
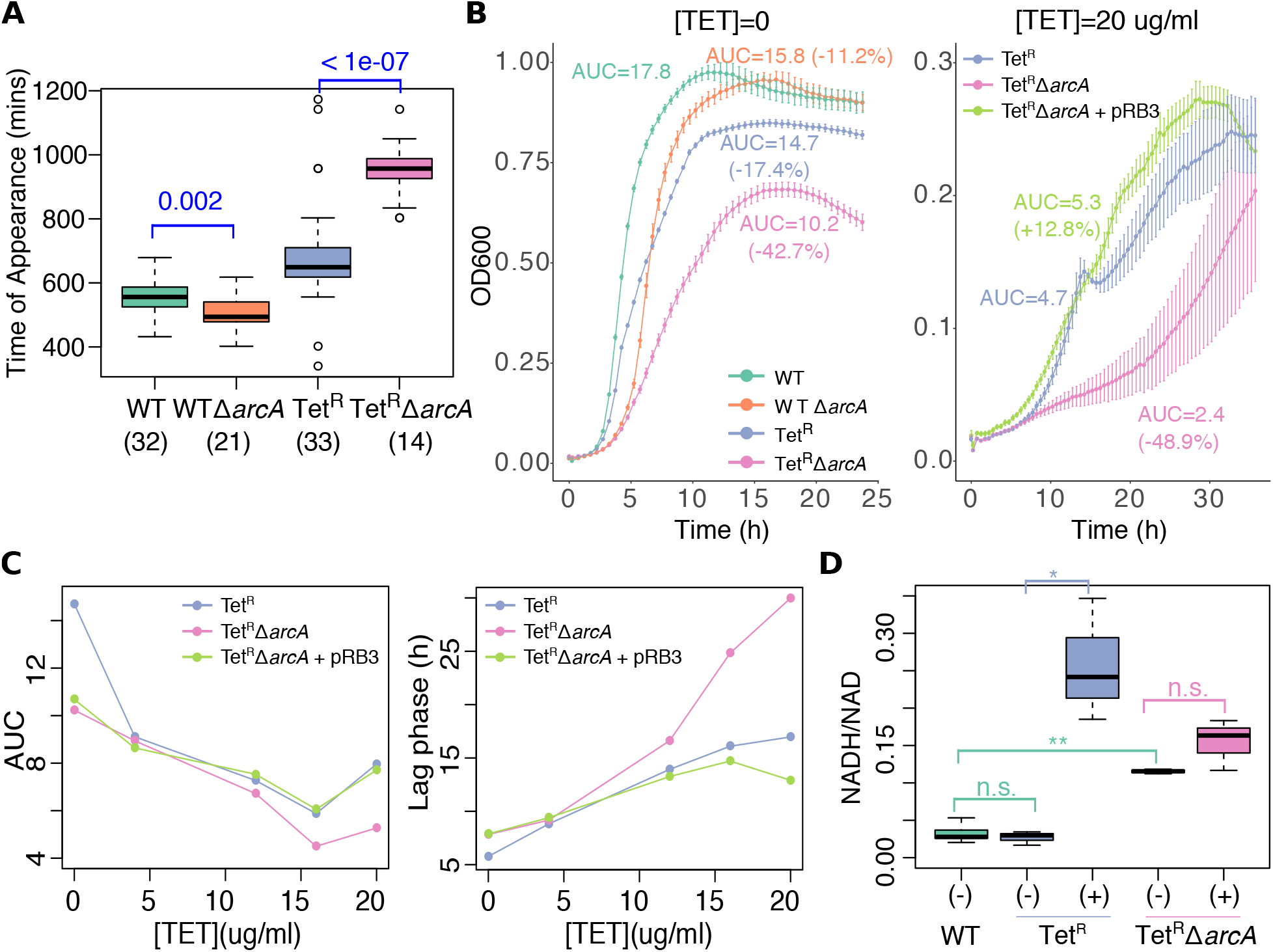
ArcA restores fitness of tetracycline resistant *E. coli*. (A) Effect of *arcA* deletion in the time of appearance, measured with the ScanLag technology^48^, of the WT and Tet^R^ strains in solid antibiotic-free LB medium. Number of colonies analyzed for each genotype are indicated in parentheses. P-values associated with Welch’s t-test for the WT vs WT Δ*arcA* and the Tet^R^ vs Tet^R^ Δ*arcA* comparisons are shown in blue. (B) Representative microdilution growth curves in LB medium at low and high tetracycline concentrations. Four replicates per strain were used. Fitness was estimated as the area under the growth curve (AUC). In the left panel, number in parentheses indicate the decrease in fitness with respect to the WT strain. In the right panel, numbers in parentheses indicate the change in fitness with respect to the Tet^R^ strain. In the Tet^R^ Δ*arcA* + pRB3 strain, the *arcA* deletion is complemented with a copy of *arcA* in the pRB3 plasmid. (C) AUC and lag phase (approximated with the Growthcurver-estimated time of inflection, the time required to achieve half of maximal OD^49^) for five different concentrations of tetracycline ([TET]). (D) NADH/NAD ratio during log phase of the WT, Tet^R^ and Tet^R^ Δ*arcA* strains. Triplicates were used for all strains/growth conditions but WT, for which six replicates were analyzed. Absence and presence of tetracycline in the growth condition is indicated with ‘(-)’ and ‘(+)’, respectively. NADH/NAD ratios were compared using Welch’s t-test. P-values in the 0.001-0.05 range and < 0.001 are indicated with the “*” and “**” symbols, respectively. N.S. indicates no significant difference.

It has been demonstrated that fast growth rate in *E. coli* causes an increase in intracellular NADH/NAD ratio^47^, which in turn triggers ArcA^52,53^ mediated repression of the TCA cycle and activation of overflow metabolism to prevent further redox imbalance^46^. Consistent with this sequence of events, we observed that tetracycline treatment significantly increased intracellular NADH/NAD ratio in the Tet^R^ strain, which explains the increased activity of ArcA with gain of tetracycline resistance. As expected, deletion of *arcA* resulted in constitutive dysregulation of NADH/NAD ratio irrespective of tetracycline treatment, presumably due to disruption of the ArcA-mediated feedback mechanism to manage redox balance (**Fig. 3D)**. We propose based on these results that ArcA plays a central role in modulating redox balance to support increased efflux-mediated tetracycline resistance phenotype.

### Drugs that mimic ArcA knockout phenotype disrupt efflux-mediated tetracycline resistance

In order to identify drugs that would simulate an *arcA* knockout phenotype, we leveraged similarities between metabolome profiles of drug-treated and single gene deletion strains of *E. coli^23^.* Among the 1,279 FDA-approved compounds in this analysis, two compounds, sertraline (a serotonin uptake inhibitor used as antidepressant) and cefpiramide (a third generation cephalosporin), generated metabolome profiles that were most similar to the metabolome of the *arcA* deletion strain (**Fig. 4A**). Reciprocally, out of ~3,800 gene deletions, *arcA* deletion was among the top 20 strains whose metabolomes were most similar to metabolome profiles generated by the two compounds. We reasoned that for the metabolome similarities to be physiologically meaningful and clinically relevant, the concentration of the drug needed to be equal to or less than the MIC for the Tet^R^ strain. While the sertraline concentration used in the metabolomics study was within its MIC, the cefpiramide concentration in the metabolic profiling study (100uM equivalent to 61.3 ug/ml)^23^ was higher than the estimated MIC (< 20 ug/ml). Hence, we excluded cefpiramide from further analysis and proceeded to test whether sertraline could resensitize Tet^R^ *E. coli* to tetracycline.

**Fig. 4.**
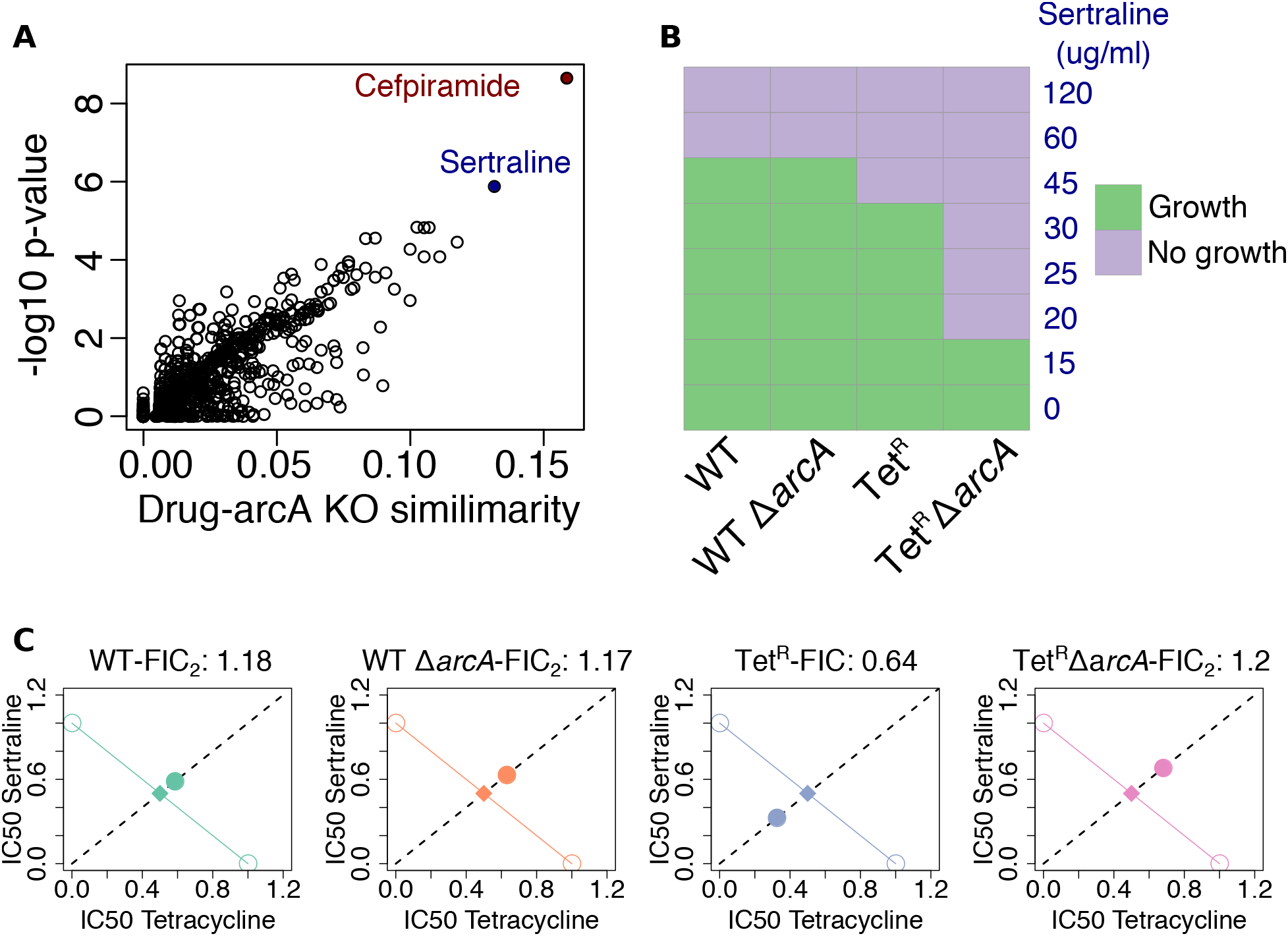
ArcA plays a central role in the synergistic effect of the sertraline-tetracycline combination in the tetracycline resistant *E. coli*. (A) Analysis of similarity between the metabolic profile of the *E. coli* treated with 1,279 FDA-approved compounds and the metabolic profile of the *arcA E. coli* deletion strain, computed by Campos and Zampieri^23^, identified sertraline (a serotonin uptake inhibitor) and cefpiramide (a third generation cephalosporin) as promising candidate ArcA inhibitors. (B) Growth assays in a microdilution series from 0 ug/ml to 120 ug/ml were performed to assess dose-response and susceptibility of the different strains to sertraline (10 replicates per strain/concentration). “Growth” indicates at least five (out of 10) replicates increased in OD by an increment of 0.1 OD after 24 hours incubation. (C) Results of DiaMOND assays to estimate fractional inhibitory concentration (FIC_2_) for the sertraline-tetracycline combinations in the four strains^54^. Five replicates per strain were used. Diamonds and filled circles indicate the expected IC50 for the sertraline-tetracycline combination in case of additive effect and the actual observed IC50 for the sertraline-tetracycline combination, respectively.

Based on similarity of metabolome profiles due to sertraline-treatment and *arcA* knockout, we predicted that the mechanism of bactericidal activity of sertraline is through its action on the ArcA network. Consistent with the increased importance of ArcA-induced response in tetracycline resistance, sertraline had a significantly greater bactericidal effect on the Tet^R^ strain, relative to the WT strain. Notably, deletion of *arcA* in the WT background had no effect on bactericidal effect of sertraline. By contrast, the bactericidal activity of sertraline was higher upon knocking out *arcA* in the Tet^R^ background (**Fig. 4B**). These results suggest that sertraline kills *E. coli* by disrupting the ArcA network, and its increased activity on the Tet^R^ *ΔarcA* strain is akin to an additive or synergistic effect due to a double hit on the same network. We performed a DiaMOND assay^54^ to test this hypothesis by investigating dose-dependent combinatorial effects of tetracycline and sertraline on WT and Tet^R^ strains with and without *arcA* deletion (**Fig. 4C**). The bactericidal activity of the tetracycline-sertraline combination was antagonistic on the WT strain, with no change upon deletion of *arcA* (fractional inhibitory concentration-FIC_2_ score ~1.2). In stark contrast, the drug combination was synergistic in killing the Tet^R^ strain (FIC_2_ score ~ 0.6), but antagonistic on the Tet^R^ Δ*arcA* strain (FIC_2_ score ~ 1.2), demonstrating unequivocally that synergy between tetracycline and sertraline emerges from disruption of the compensatory physiologic state that is mechanistically generated by ArcA activity.

## DISCUSSION

We have discovered that global remodeling of transcription by a network of at least 25 TFs generates a novel metabolic state to compensate for the gain of tetracycline resistance in *E. coli.* Interestingly, while the resistance mutations resulted in constitutive overexpression of the *acrAB* operon in the Tet^R^ strain, the global transcriptional remodeling manifested in a dramatic manner only during tetracycline treatment, demonstrating that it was a downstream consequence of the increased activity of the AcrAB efflux pump. We propose a model to explain how increased efflux triggers a compensatory physiologic state to support tetracycline resistance in Tet^R^ *E. coli* (**Fig 5**). AcrAB is an efflux pump of the RND (Resistance-Nodulation-Division) superfamily that consumes the proton motive force (PMF) to expel intracellular substrates, tetracycline in this case^55^. Hence, AcrAB competes with the ETC and ATP synthase for both space in the membrane as well as the PMF, which is required for ATP synthesis^46^. This competition may reduce oxidation of NADH molecules by the ETC, and the resulting increase in NADH/NAD ratio triggers ArcA. ArcA acts by repressing the TCA cycle and redirecting metabolic flux away from oxidative phosphorylation towards overflow metabolism as an alternative sources of energy^45,47,52,53^. This hypothesis is supported by two independent observations: (i) deleting either of the two repressors (*marR* and *acrR*) of the *acrAB* operon resulted in the increased secretion of acetate, a byproduct of overflow metabolism^35^; and (ii) mutations in *acrB* significantly reduced the rate of oxygen uptake^35^. The downregulation of the TCA cycle may also serve to mitigate oxidative stress by preventing the production of reactive oxygen species (ROS)^56^, although that is unlikely to be the case here since bacteriostatic antibiotics such as tetracycline and chloramphenicol do not increase the production of ROS^57,58^. The global remodeling of the respiration and energy production pathways appears to be a generalized mechanism for restoring fitness across AMR pathogens, including *Pseudomonas aeruginosa^17^* and *Chromobacterium violaceum^22^*, that also show high intracellular NADH levels upon gaining resistance to diverse antibiotics through the increased expression of RND efflux pumps.

**Fig. 5.**
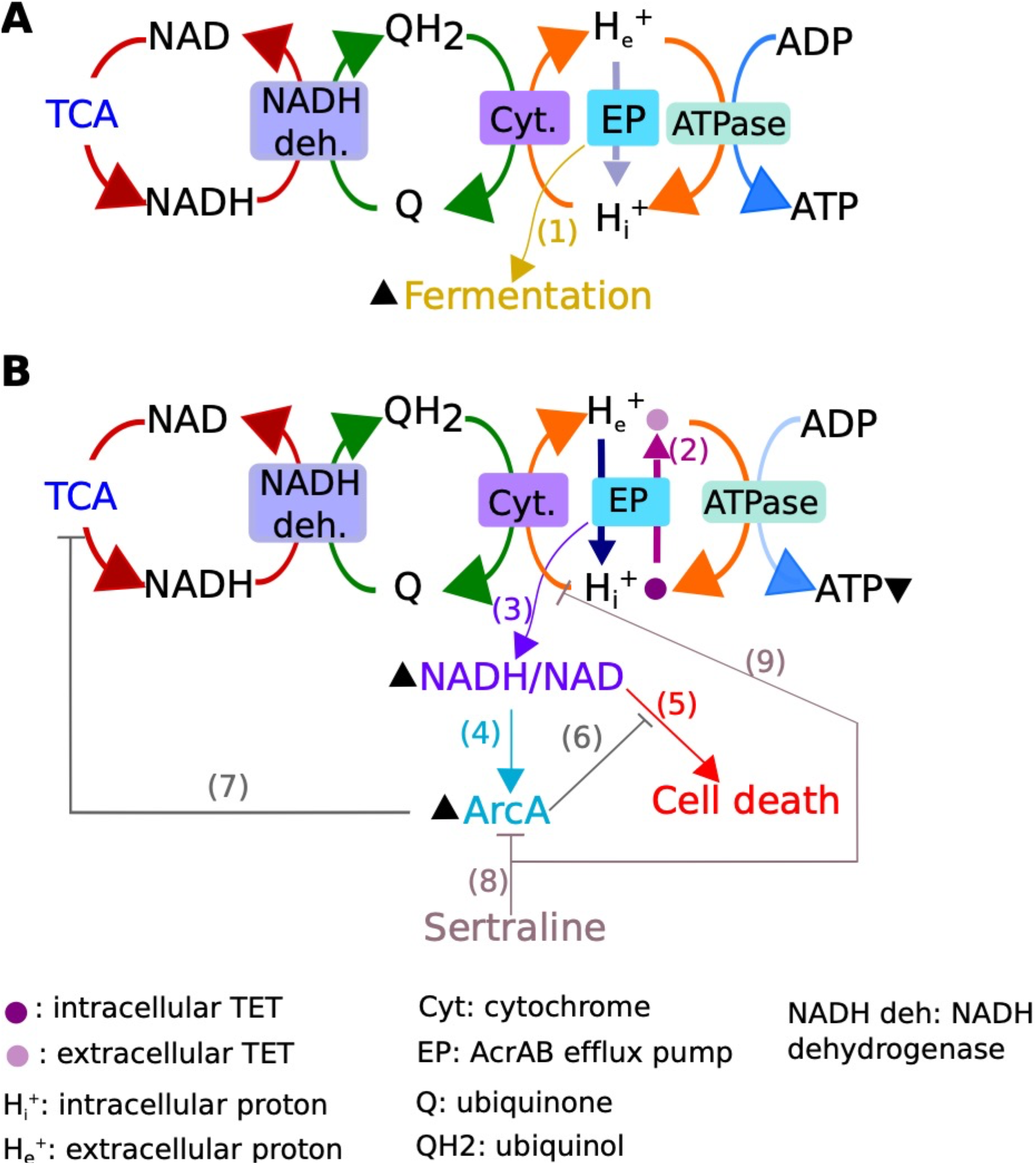
Summary overview of the ArcA-driven compensatory mechanism for tetracycline resistance and its connection with efflux pump mediated resistance. (A) In the absence of tetracycline, Tet^R^ upregulates the *acrAB* efflux pump (EP), which in turn causes upregulation of fermentation-related genes (edge #1). This observation is independently supported by excretion of fermentation byproducts when transcriptional repressors of the AcrAB pump are deleted^35^. (B) The activity of AcrAB increases in the presence of tetracycline^28^ (edge #2), driving dramatic changes in metabolism. We hypothesize that AcrAB disrupts the functions of other transporters by crowding the membrane, consuming the proton motive force, causing an increase in NADH/NAD ratio (edge #3, **Fig. 3D**), which activates ArcA^52^ (edge #4, **Fig. 2A**). The potential toxic consequence of increased NADH/NAD ratio^46^ (edge #5), is alleviated by ArcA (edge #6) through downregulation of the TCA cycle^44^ (edge #7). Finally, sertraline represses the ArcA network (edge #8, **Fig. 4A**) and putatively inhibits the PMF^61^ (edge #9) to synergistically potentiate the bacteriostatic effect of tetracycline. Relative increase in abundance or activity are indicated with an upward pointing arrowhead, next to the corresponding molecule. Decreased concentration is indicated with a downward pointing arrowhead.

Amplification of the fitness cost of tetracycline resistance upon disrupting the master regulator (ArcA) of the compensatory metabolic state demonstrated how a network-based approach can rationally identify new vulnerabilities that emerge as a consequence of gaining resistance, because knocking out ArcA had a minor fitness consequence in the WT background (**Fig. 2** and **Fig. 3**). Having identified ArcA as a new vulnerability in the Tet^R^ strain, we were able to identify secondary molecule(s) that could target the compensatory physiologic state by leveraging the similarities between global metabolome changes in *E. coli* across a library of single gene deletion strains, and a drug library screen^23^. This analysis rank-prioritized the most likely drugs in the screen that could disrupt the ArcA network to potentiate tetracycline action. Sertraline, which was among the top ranking candidates, was proposed to potentiate tetracycline action by blocking the PMF and indirectly inhibiting its efflux^59–61^, albeit by a different plasmid-encoded TetA pump^25,61^. The loss of synergistic action of tetracycline-sertraline combination upon deleting *arcA* demonstrated the mechanism of synergy (**Fig. 4C**), but also revealed how Tet^R^ *E. coli* could (re)gain resistance and tolerance to this drug combination through a single regulatory mutation or transcriptional reprogramming of a single TF. Notably, the network analysis in this study implicated at least 25 TFs and their networks might directly contribute towards the compensatory physiologic state required to support the tetracycline resistance phenotype of Tet^R^. The complexity of this network illustrates the multiple routes through which a pathogen like *E. coli* could escape antibiotic treatment to gain resistance, explaining why we need multi-drug combinations to combat antibiotic tolerance and resistance^62–64^.

Formulating multi-drug regimen is particularly challenging because the numbers of combinations that need to be tested is too large, even for a high throughput drug screen^65^. The network-based approach described in this study will prove valuable in this effort because it uses a mechanistic model of a gene regulatory network underlying tolerance^62^ or resistance phenotypes to rank prioritize combinations of molecules that target multiple vulnerabilities within a pathogen. We have demonstrated that this strategy could enable the recovery of “lost” antibiotics by identifying new vulnerabilities that emerge within transcriptional and metabolic networks to manage tradeoff in fitness upon gain of resistance^3,17^. This approach can be combined with laboratory evolution experiments to delineate trajectories of antibiotic resistance^66^ and design drug combinations to preemptively curtail emergence and spread of AMR. For instance, antibiotic tolerant strains of *E. coli* gained resistance at a faster rate than the wild type strain, suggesting that drugs that target the tolerance networks within these strains could potentially delay or block the emergence of resistance^66^. Another high throughput screen of a library of TF deletion strains demonstrated that deletion of *arcA* suppresses gain of resistance to cefixime, ciprofloxacin and chloramphenicol^56^. Based on this observation, we posit that multi-drug regimen formulated based on vulnerabilities within networks governed by ArcA and other TFs identified in this study could have generalized value in extending the lifespans of a broad range of existing antibiotics, and supplement the development of new antimicrobial compounds^5^.

## METHODS

### *E. coli* strains and culturing conditions

*E. coli* MG1655 (WT) and a lab-evolved tetracycline resistant *E. coli* derived from the susceptible WT (Tet^R^) were kindly provided by Professor Benno ter Kuile^30^. *arcA* mutants were generated using the Red^®^/ET^®^ recombination kit (Gene Brigdes, Germany), following the manufacturers’ instructions. Using this approach, a kanamycin cassette was inserted between the 5’ and 3’ regions (50bp) of *arcA. arcA* disruption was confirmed with PCR and Sanger sequencing. *arcA* deletion was complemented with the pRB3-*arcA* plasmid^67^ kindly provided by Professor Sangwei Lu. *E. coli* strains were grown on Luria-Beltrani (LB) on agar and broth (with constant shaking) at 37°C and aerobic conditions.

### Assessment of *arcA* deletion effect on fitness

Microdilution growth curves were performed in LB broth using a Bioscreen C instrument (Growth Curves USA, Piscataway, NJ). First, frozen cells were used to start overnight cultures that were then used as inoculum for the growth assays. WT and WT ΔarcA precultures were prepared in tetracycline-free medium. The Tet^R^, Tet^R^ ΔarcA and Tet^R^ *arcA*^+^ (complemented strain) precultures contained 4 ug/ml of tetracycline (Sigma-Aldrich) to maintain selection pressure for resistance. Overnight cultures were adjusted to 0.01 OD and 200 uL cultures were run at 37°C and continuous shaking. OD was measured every 30 mins. Wells only inoculated with LB were included to measure background optical density of the medium. The Growthcurver R package^49^ was used to fit a logistic equation to the OD data after blank correction. Bacterial fitness was estimated in terms of the area under the growth curve (AUC) empirically determined by Growthcurver. The AUC value integrates multiple properties of the growth curve^49^. For this reason and consistency among biological replicates^50^, it has been recently preferred over other fitness-related terms^50,51^. To evaluate fitness in solid LB medium (incubated at 37°C), the time required to detect colonies in petri dishes (time of appearance) was determined using the ScanLag technology^48^ for the analyzed strains. Results were consistent among different pixel thresholds (to define when a colony was considered visible). Petri dishes were inoculated with the 10^-4^ dilution of the overnight cultures previously adjusted to 0.01OD.

### DiaMOND assay to evaluate synergy of two drug combinations

We used the diagonal measurement of n-way drug interactions (DiaMOND) assay^54^ (as described by Cokol-Cakmak et al.^68^) to evaluate the predicted ArcA-mediated synergy between tetracycline and selected compounds (sertraline hydrochloride, Sigma-Aldrich). Briefly, we first determined the single drug concentrations that reduced OD (measured after 16 hours of growth) by half with respect to the drug-free condition (corresponding to the drug IC50), using a BioTek Epoch 2 instrument (BioTek, USA). Spline fitting was used to interpolate the IC50 values in the analyzed growth curves. Then, the IC50 of the two-drug combinations (between tetracycline and selected compounds in a 1:1 volume using the IC50 concentrations defined in the previous step) were determined. The fractional inhibitory concentration for the two-drug combinations (FIC_2_) under the Loewe additivity model was defined^54^. Cultures inoculum were prepared as described before for the fitness assays with the distinction that all overnight cultures were performed in drug-free medium to avoid any cross-reaction with the other tested compounds.

### NADH/NAD ratio measurements

NADH and NAD ratios were measured using the Enzychrom NAD/NADH assay kit (Bioassay Systems), following the manufacters’ instructions (and adding a sonication step of 20 seconds, before heating at 60°C for 5 mins to lyse the bacterial cells). To simultaneously capture in log phase all cultures, overnight cultures were adjusted to slightly different ODs to correct for differences in fitness among strains (i.e. the inoculum with the highest OD was the one corresponding to the Tet^R^ ΔarcA strain while the inoculum with the lowest OD was the one corresponding to the WT strain). In each experiment, the NADH/NAD ratio of the WT strain (without tetracycline) was measured to evaluate consistency between experiments (that included three replicates per strain).

### Differential expression analysis of *E. coli* MG1655 and tetracycline resistant MG1655 microarray data

To characterize the adaptation of the Tet^R^ strain to tetracycline, we analyzed the publicly available normalized microarray data for the WT and Tet^R^ strains in the presence and absence of tetracycline (GEO accession no. GSE57084) reported by Händel et al^30^. Differential expression analysis of microarray data was performed using a Bayesian T-test with the Cyber-T tool^69^. Genes with adjusted p-values < 0.05 and absolute log2 fold-change >1 were considered differentially expressed.

### Functional enrichment analysis

Enrichment analyses of significantly up- and down-regulated set of genes were independently performed using DAVID^70^. Only functional terms with adjusted p-values (Benjamini-Hochberg) < 0.05 were considered enriched.

### Identification of differentially active regulatory circuits associated with the gain of tetracycline resistance

We identified differentially active TFs using the NetSurgeon algorithm^38^, as previously applied for *Mycobacterium tuberculosis*^71^. Briefly, NetSurgeon ranks TFs based on their potential influence in the observed transcriptional changes between two states of interest (estimated according to the change in expression of their known target genes). 192 TF regulons were extracted from a transcriptional network (containing 5,517 signed TF-gene interactions) compiled from the RegulonDB version 9.0 database^37^. 68 out of the 192 TFs had less than five target genes and were not included in the analysis to reduce false positives due to overlap between regulons. We focused on the transcriptional changes between the Tet^R^ strain and the parental WT strain in the absence of tetracycline, and the response of the Tet^R^ strain to tetracycline. TFs ranked (using the highest score between the independently computed scores for increased activity and decreased activity) in the top 15 of each comparison were considered as differentially active. To complement the NetSurgeon analysis, a network component analysis^40^ was applied to estimate the TFs activity (TFA) using the transcriptional profile of their known targets. The RegulonDB-derived transcriptional network described above and the microarray data reported by Händel et al. were used to estimate the TFA as previously described^30,72^. Statistical differences in TF activity were determined using a Welch’s T-test. Only TFs with p-values < 0.05 were considered differentially active. Finally, we mined the EGRIN model previously developed for *E. coli*^41^ to identify clusters of co-regulated genes statistically enriched (with adjusted hypergeometric test p-values < 0.05 and containing >9 DEGs) with genes whose expression was altered by gain of tetracycline resistance. The association between clusters with differentially expressed genes and TFs was used as an indicator of differential activity of the relevant TFs.

## Acknowledgements

We thank members of the Baliga lab for critical discussions and feedback. We thank Professor Benno ter Kuile at University of Amsterdam for kindly providing the *E. coli* strains used in this study. We also thank Professor Sangwei Lu at University of California Berkeley for kindly providing the pRB3-*arcA* plasmid. Funding was provided by the National Science Foundation (DBI-1565166), NIH (R01AI141953 and R01AI128215) to NSB.

## SUPPLEMENTARY FIGURES

**Fig S1.**
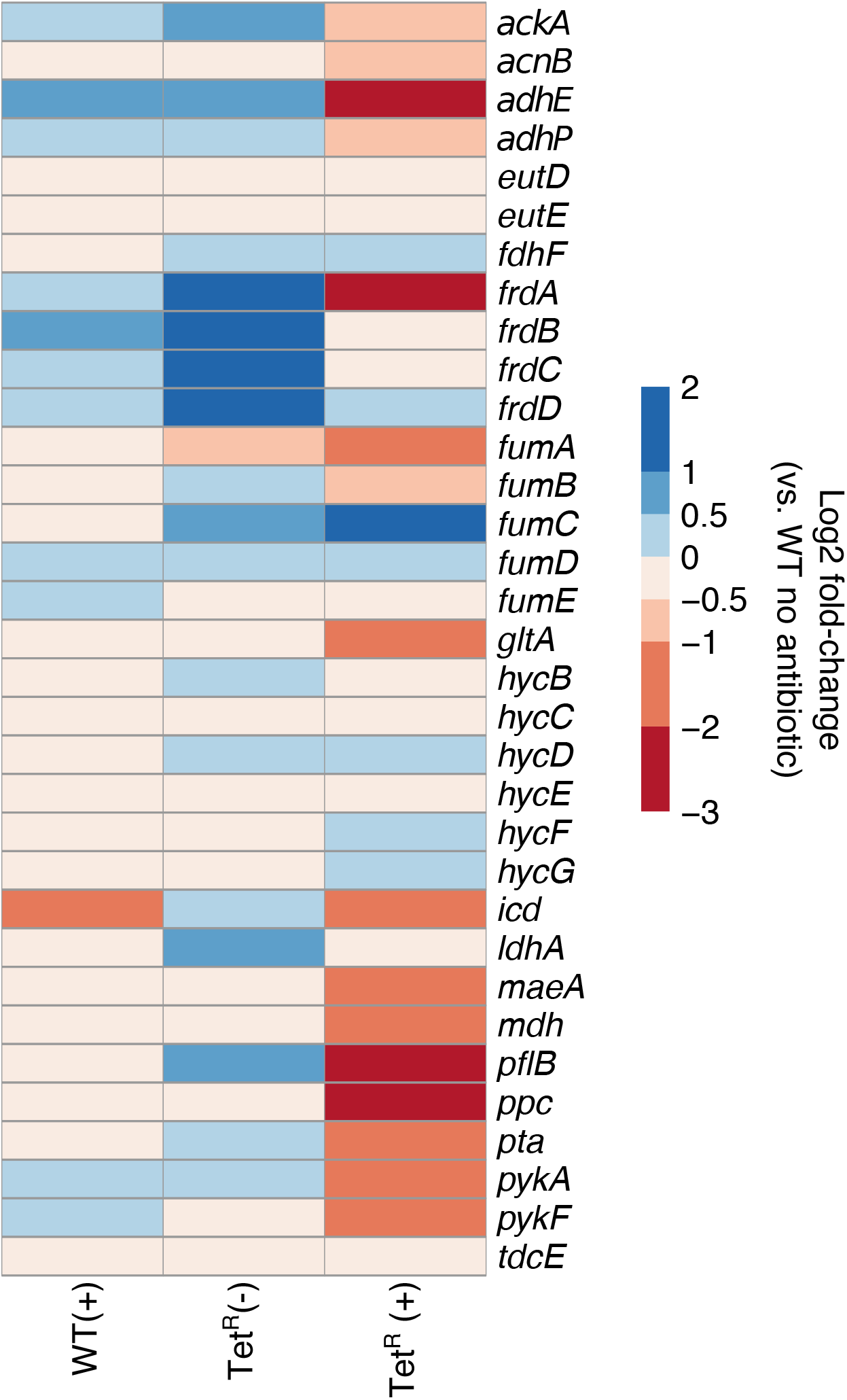
Relative transcript level changes of fermentation-related genes in the Tet^R^ strain. The heat map shows relative log_2_ fold-change in transcript levels of fermentation-related genes (compiled from the EcoCyc database^73^) in the WT and Tet^R^ strain with (+) and without (-) tetracycline treatment. All fold change values are relative to transcript levels in the WT strain without tetracycline treatment.

**Fig S2.**
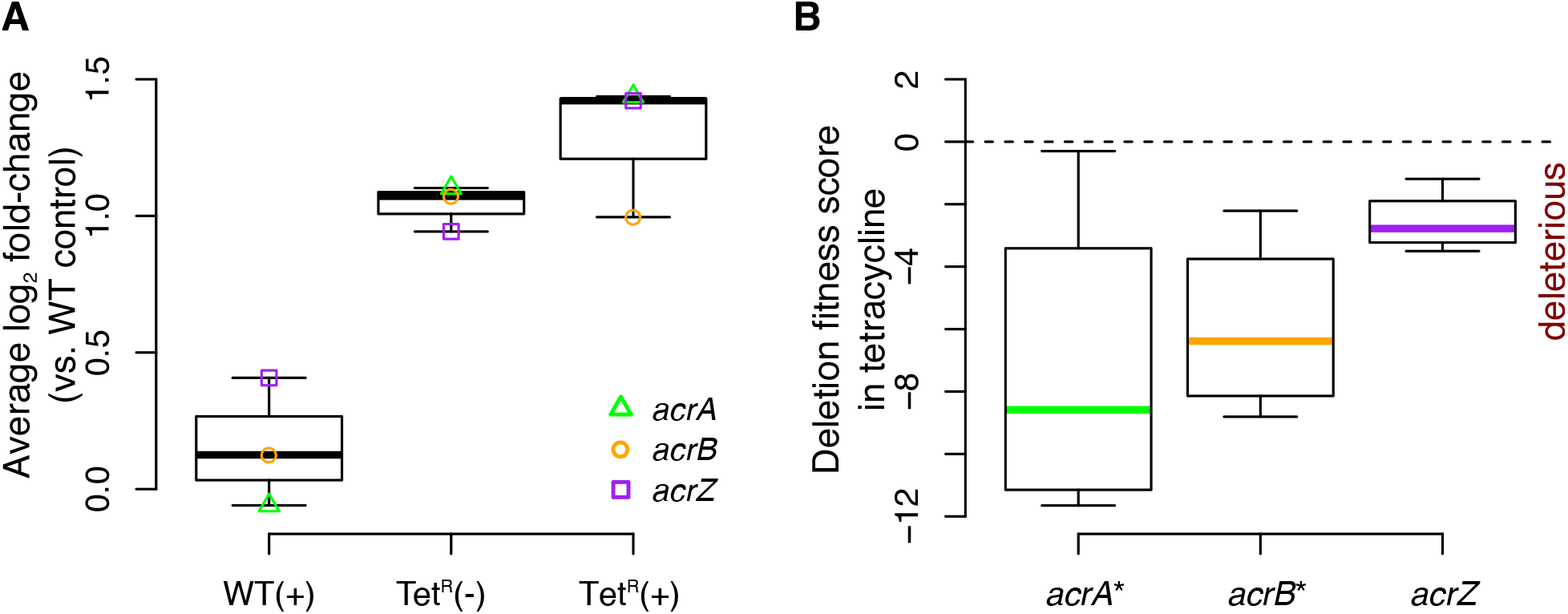
The AcrAB efflux pump contributes to tetracycline resistance. (A) The *acrAB* operon and *acrZ* (that encodes a protein that interacts with the AcrAB pump and confers resistance to tetracycline^31^) were constitutively up-regulated in the Tet^R^ strain^30^. (B) Fitness effect of single deletions of the *acrAB* and *acrZ* during tetracycline treatment in the WT genotype (from Nichols et al^33^). The “*” symbol indicates that deletion of *acrA* and *acrB* were among the top 10 most deleterious effects on tetracycline resistance phenotype across the library of ~3,800 gene knockouts.

**Fig S3.**
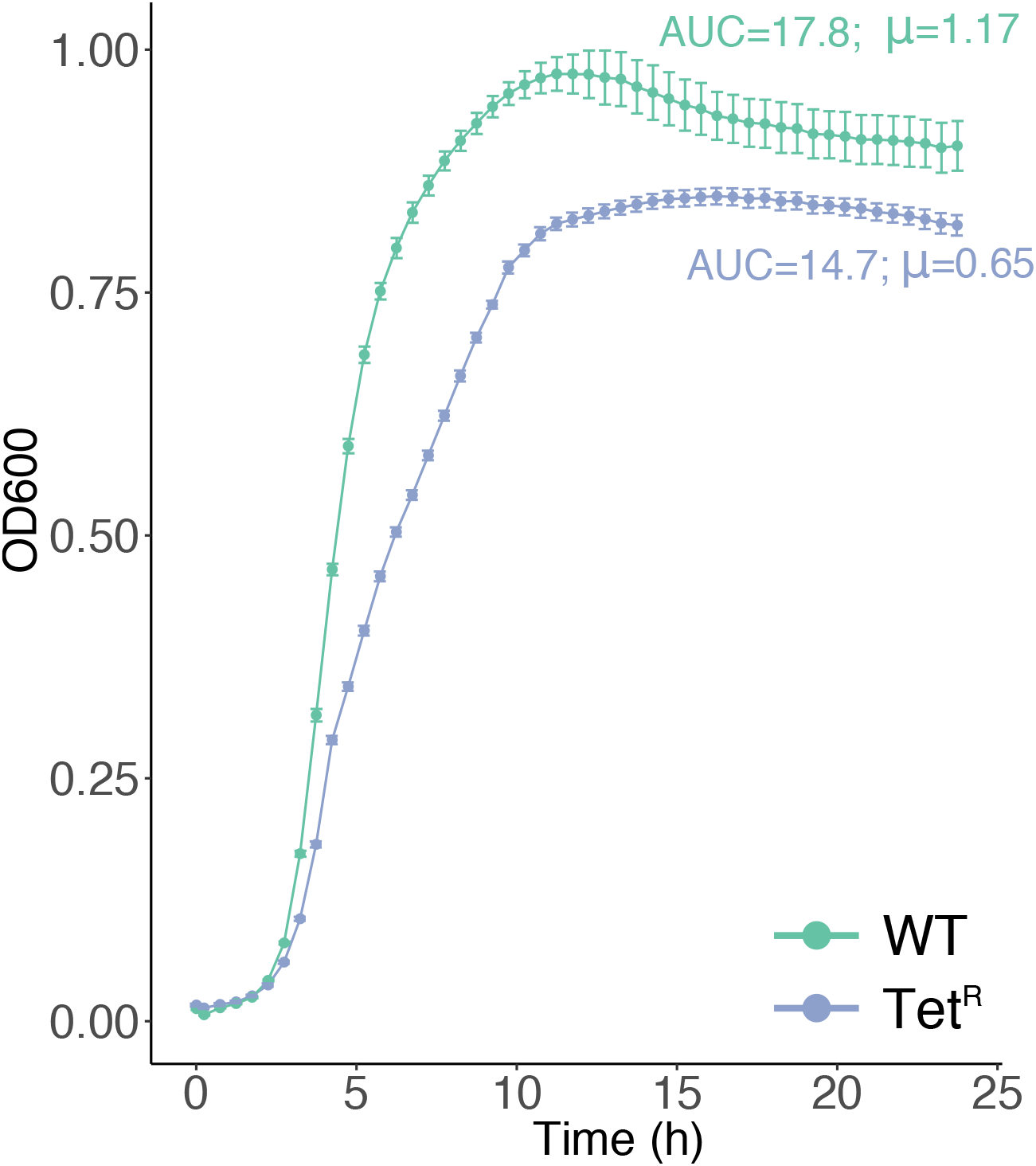
Fitness cost associated with the gain of tetracycline resistance. Growth curves of four replicate cultures of WT and Tet^R^ strains in LB medium without tetracycline. Fitness was estimated as the area under the growth curve (AUC) using the Growthcurver R package^49^. The average Growthcurver-derived maximum growth rates (“μ”) are also shown.

## SUPPLEMENTARY TABLES

**Table S1.**
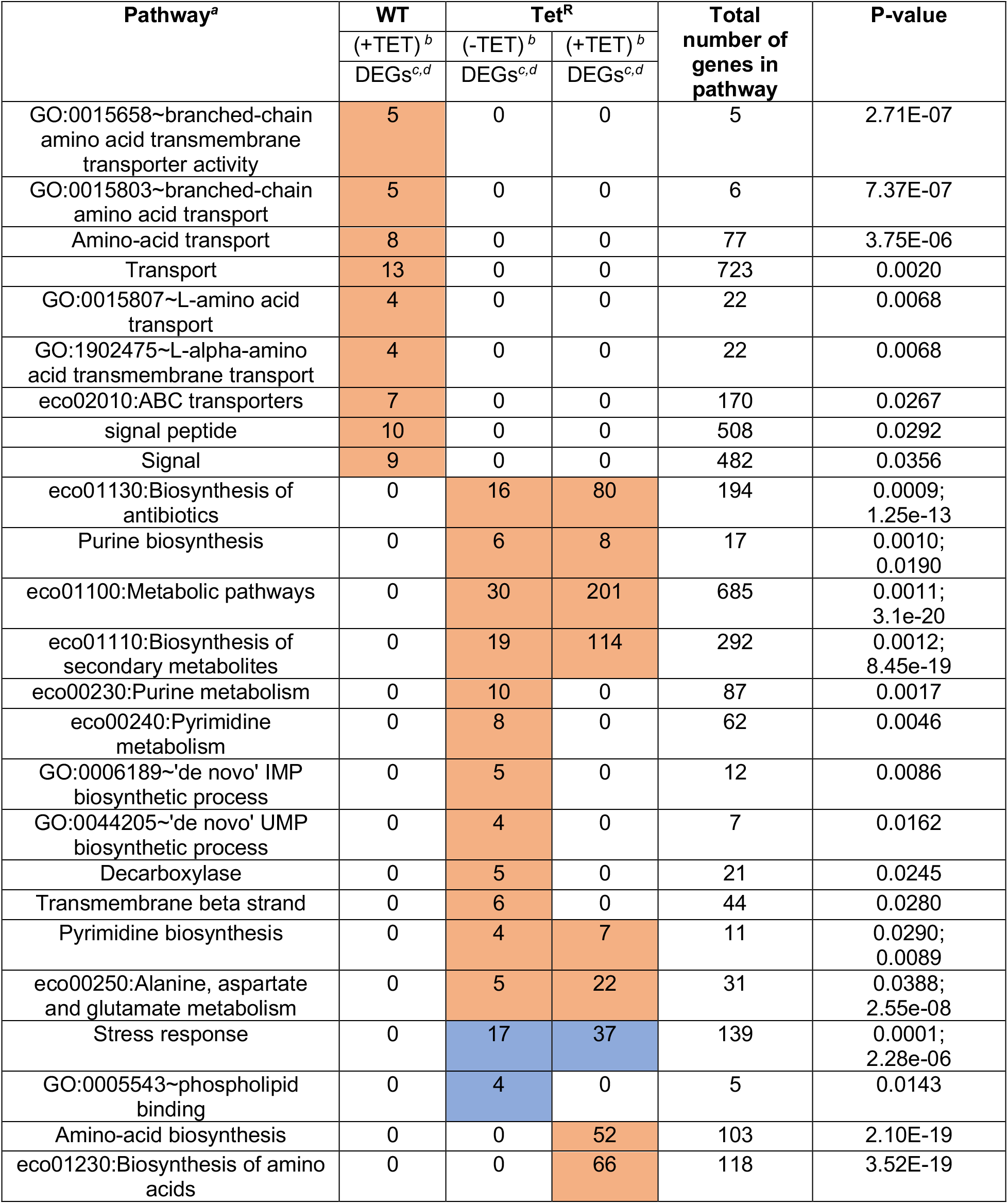
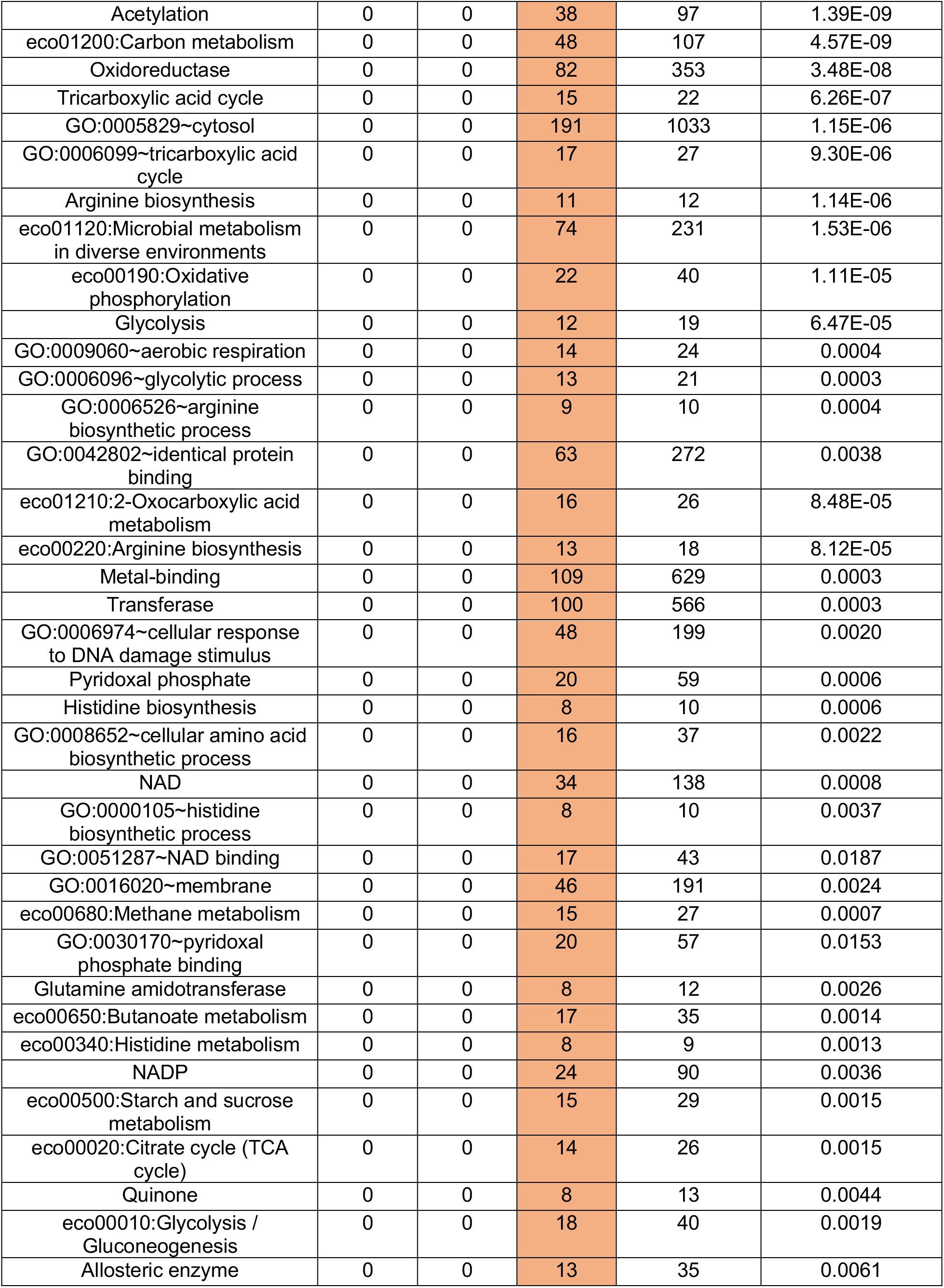
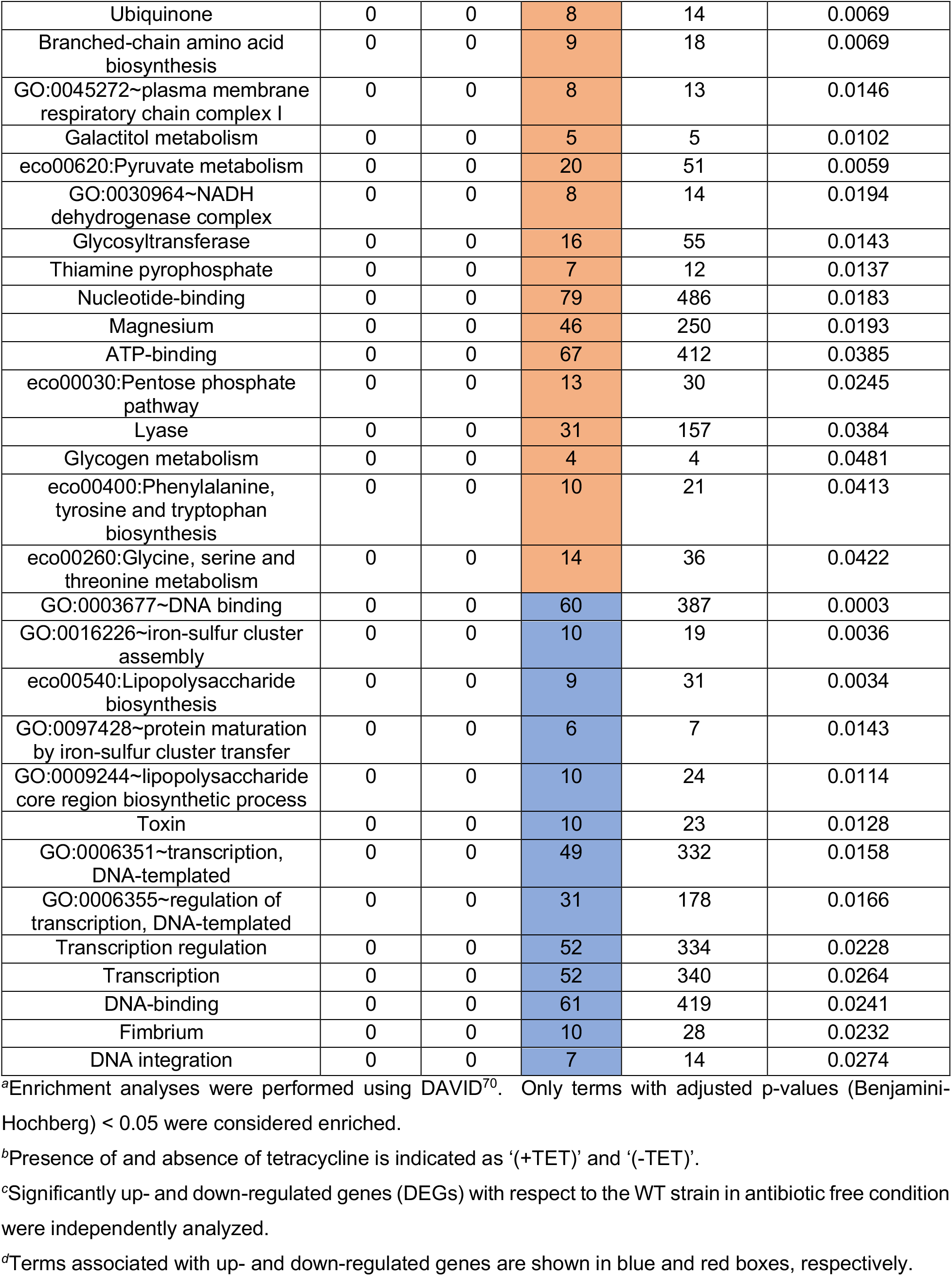
Functional enrichment information of genes differentially expressed in the WT and Tet^R^ strains.

## REFERENCES

1. Blair, J. M. A., Webber, M. A., Baylay, A. J., Ogbolu, D. O. & Piddock, L. J. V. Molecular mechanisms of antibiotic resistance. Nat. Rev. Microbiol. 13, 42–51 (2015).

2. Brauner, A., Fridman, O., Gefen, O. & Balaban, N. Q. Distinguishing between resistance, tolerance and persistence to antibiotic treatment. Nat. Rev. Microbiol. 14, 320–330 (2016).

3. Andersson, D. I. & Hughes, D. Antibiotic resistance and its cost: is it possible to reverse resistance? Nat. Rev. Microbiol. 8, 260 (2010).

4. O’Neill, J. Tackling drug-resistant infections globally: final report and recommendations. (Review on Antimicrobial Resistance, 2016).

5. Ventola, C. L. The antibiotic resistance crisis: part 1: causes and threats. Pharm. Ther. 40, 277 (2015).

6. Zhang, Q.-Q., Ying, G.-G., Pan, C.-G., Liu, Y.-S. & Zhao, J.-L. Comprehensive evaluation of antibiotics emission and fate in the river basins of China: source analysis, multimedia modeling, and linkage to bacterial resistance. Environ. Sci. Technol. 49, 6772–6782 (2015).

7. Kohanski, M. A., DePristo, M. A. & Collins, J. J. Sublethal antibiotic treatment leads to multidrug resistance via radical-induced mutagenesis. Mol. Cell 37, 311–320 (2010).

8. Long, H. et al. Antibiotic treatment enhances the genome-wide mutation rate of target cells. Proc. Natl. Acad. Sci. 113, E2498–E2505 (2016).

9. Kollef, M. H. & Fraser, V. J. Antibiotic resistance in the intensive care unit. Ann. Intern. Med. 134, 298–314 (2001).

10. Mölstad, S. et al. Sustained reduction of antibiotic use and low bacterial resistance: 10-year follow-up of the Swedish Strama programme. Lancet Infect. Dis. 8, 125–132 (2008).

11. Weis, S. E. et al. The effect of directly observed therapy on the rates of drug resistance and relapse in tuberculosis. N. Engl. J. Med. 330, 1179–1184 (1994).

12. Cars, O., Mölstad, S. & Melander, A. Variation in antibiotic use in the European Union. Lancet 357, 1851–1853 (2001).

13. Fernandes, P. & Martens, E. Antibiotics in late clinical development. Biochem. Pharmacol. 133, 152–163 (2017).

14. Lewis, K. Platforms for antibiotic discovery. Nat. Rev. Drug Discov. 12, 371–387 (2013).

15. Suzuki, S., Horinouchi, T. & Furusawa, C. Prediction of antibiotic resistance by gene expression profiles. Nat Commun 5, 5792 (2014).

16. Dunphy, L. J., Yen, P. & Papin, J. A. Integrated Experimental and Computational analyses reveal differential metabolic functionality in antibiotic-resistant *Pseudomonas aeruginosa*. Cell Syst. 8, 3–14 (2019).

17. Pacheco, J. O., Alvarez-Ortega, C., Rico, M. A. & Martinez, J. L. Metabolic compensation of fitness costs is a general outcome for antibiotic-resistant *Pseudomonas aeruginosa* mutants overexpressing efflux pumps. MBio 8, e00500–17 (2017).

18. Freihofer, P. et al. Nonmutational compensation of the fitness cost of antibiotic resistance in mycobacteria by overexpression of tlyA rRNA methylase. Rna 22, 1836–1843 (2016).

19. Ejim, L. et al. Combinations of antibiotics and nonantibiotic drugs enhance antimicrobial efficacy. Nat. Chem. Biol. 7, 348 (2011).

20. Baym, M., Stone, L. K. & Kishony, R. Multidrug evolutionary strategies to reverse antibiotic resistance. Science (80-.). 351, (2016).

21. Peng, B. O. et al. Exogenous alanine and/or glucose plus kanamycin kills antibiotic-resistant bacteria. Cell Metab. 21, 249–262 (2015).

22. Banerjee, D. et al. A scalable metabolite supplementation strategy against antibiotic resistant pathogen *Chromobacterium violaceum* induced by NAD+/NADH+ imbalance. BMC Syst. Biol. 11, 1–20 (2017).

23. Campos, A. I. & Zampieri, M. Metabolomics-driven exploration of the chemical drug space to predict combination antimicrobial therapies. Mol. Cell 74, 1291–1303 (2019).

24. Roberts, M. C. Tetracycline resistance determinants: mechanisms of action, regulation of expression, genetic mobility, and distribution. FEMS Microbiol. Rev. 19, 1–24 (1996).

25. Thaker, M., Spanogiannopoulos, P. & Wright, G. D. The tetracycline resistome. Cell. Mol. Life Sci. 67, 419–431 (2010).

26. FDA, U. S. Antimicrobials sold or distributed for use in food-producing animals. (2017).

27. Nelson, M. L. & Levy, S. B. Reversal of tetracycline resistance mediated by different bacterial tetracycline resistance determinants by an inhibitor of the Tet (B) antiport protein. Antimicrob. Agents Chemother. 43, 1719–1724 (1999).

28. Viveiros, M. et al. Inducement and reversal of tetracycline resistance in *Escherichia coli* K-12 and expression of proton gradient-dependent multidrug efflux pump genes. Antimicrob. Agents Chemother. 49, 3578–3582 (2005).

29. van der Horst, M. A., Schuurmans, J. M., Smid, M. C., Koenders, B. B. & ter Kuile, B. H. De novo acquisition of resistance to three antibiotics by *Escherichia coli*. Microb. drug Resist. 17, 141–147 (2011).

30. Händel, N., Schuurmans, J. M., Feng, Y., Brul, S. & Ter Kuile, B. H. Interaction between mutations and regulation of gene expression during development of de novo antibiotic resistance. Antimicrob. Agents Chemother. 58, 4371–4379 (2014).

31. Hobbs, E. C., Yin, X., Paul, B. J., Astarita, J. L. & Storz, G. Conserved small protein associates with the multidrug efflux pump AcrB and differentially affects antibiotic resistance. Proc. Natl. Acad. Sci. 109, 16696–16701 (2012).

32. de Cristobal, R. E., Vincent, P. A. & Salomon, R. A. Multidrug resistance pump AcrAB-TolC is required for high-level, Tet (A)-mediated tetracycline resistance in *Escherichia coli*. J. Antimicrob. Chemother. 58, 31–36 (2006).

33. Nichols, R. J. et al. Phenotypic landscape of a bacterial cell. Cell 144, 143–156 (2011).

34. Hoeksema, M., Jonker, M. J., Brul, S. & Ter Kuile, B. H. Effects of a previously selected antibiotic resistance on mutations acquired during development of a second resistance in *Escherichia coli*. BMC Genomics 20, 284 (2019).

35. Zampieri, M. et al. Metabolic constraints on the evolution of antibiotic resistance. Mol. Syst. Biol. 13, 917 (2017).

36. Lázár, V. et al. Genome-wide analysis captures the determinants of the antibiotic cross-resistance interaction network. Nat. Commun. 5, 4352 (2014).

37. Gama-Castro, S. et al. RegulonDB version 9.0: high-level integration of gene regulation, coexpression, motif clustering and beyond. Nucleic Acids Res. 44, D133–43 (2016).

38. Michael, D. G. et al. Model-based transcriptome engineering promotes a fermentative transcriptional state in yeast. Proc. Natl. Acad. Sci. 113, E7428–E7437 (2016).

39. Duval, V. & Lister, I. M. MarA, SoxS and Rob of *Escherichia coli*--Global regulators of multidrug resistance, virulence and stress response. Int. J. Biotechnol. wellness Ind. 2, 101 (2013).

40. Liao, J. C. et al. Network component analysis: reconstruction of regulatory signals in biological systems. Proc. Natl. Acad. Sci. U. S. A. 100, 15522–7 (2003).

41. Brooks, A. N. et al. A system-level model for the microbial regulatory genome. Mol. Syst. Biol. 10, 740 (2014).

42. Shalel-Levanon, S., San, K.-Y. & Bennett, G. N. Effect of oxygen, and ArcA and FNR regulators on the expression of genes related to the electron transfer chain and the TCA cycle in *Escherichia coli*. Metab. Eng. 7, 364–374 (2005).

43. Covert, M. W., Knight, E. M., Reed, J. L., Herrgard, M. J. & Palsson, B. O. Integrating high-throughput and computational data elucidates bacterial networks. Nature 429, 92–96 (2004).

44. Park, D. M., Akhtar, M. S., Ansari, A. Z., Landick, R. & Kiley, P. J. The bacterial response regulator ArcA uses a diverse binding site architecture to regulate carbon oxidation globally. PLoS Genet. 9, (2013).

45. Basan, M., Hui, S. & Williamson, J. R. ArcA overexpression induces fermentation and results in enhanced growth rates of *E. coli*. Sci. Rep. 7, 1–7 (2017).

46. Szenk, M., Dill, K. A. & de Graff, A. M. R. Why do fast-growing bacteria enter overflow metabolism? Testing the membrane real estate hypothesis. Cell Syst. 5, 95–104 (2017).

47. Basan, M. et al. Overflow metabolism in *Escherichia coli* results from efficient proteome allocation. Nature 528, 99–104 (2015).

48. Levin-Reisman, I. et al. Automated imaging with ScanLag reveals previously undetectable bacterial growth phenotypes. Nat. Methods 7, 737–739 (2010).

49. Sprouffske, K. & Wagner, A. Growthcurver: an R package for obtaining interpretable metrics from microbial growth curves. BMC Bioinformatics 17, 172 (2016).

50. Palmer, A. C. et al. Delayed commitment to evolutionary fate in antibiotic resistance fitness landscapes. Nat. Commun. 6, 1–8 (2015).

51. Dunai, A. et al. Rapid decline of bacterial drug-resistance in an antibiotic-free environment through phenotypic reversion. Elife 8, (2019).

52. Vemuri, G. N., Altman, E., Sangurdekar, D. P., Khodursky, A. B. & Eiteman, M. A. Overflow metabolism in *Escherichia* coli during steady-state growth: transcriptional regulation and effect of the redox ratio. Appl. Environ. Microbiol. 72, 3653–3661 (2006).

53. Vemuri, G. N., Eiteman, M. A. & Altman, E. Increased recombinant protein production in *Escherichia coli* strains with overexpressed water-forming NADH oxidase and a deleted ArcA regulatory protein. Biotechnol. Bioeng. 94, 538–542 (2006).

54. Cokol, M., Kuru, N., Bicak, E., Larkins-Ford, J. & Aldridge, B. B. Efficient measurement and factorization of high-order drug interactions in *Mycobacterium tuberculosis*. Sci. Adv. 3, e1701881 (2017).

55. Anes, J., McCusker, M. P., Fanning, S. & Martins, M. The ins and outs of RND efflux pumps in *Escherichia coli*. Front. Microbiol. 6, 587 (2015).

56. Horinouchi, T., Maeda, T., Kotani, H. & Furusawa, C. Suppression of antibiotic resistance evolution by single-gene deletion. Sci. Rep. 10, 1–9 (2020).

57. Kohanski, M. A., Dwyer, D. J., Hayete, B., Lawrence, C. A. & Collins, J. J. A common mechanism of cellular death induced by bactericidal antibiotics. Cell 130, 797–810 (2007).

58. Hao, Z. et al. The multiple antibiotic resistance regulator MarR is a copper sensor in *Escherichia coli*. Nat. Chem. Biol. 10, 21–28 (2014).

59. Munoz-Bellido, J. L., Munoz-Criado, S. & Garcia-Rodriguez, J. A. Antimicrobial activity of psychotropic drugs: selective serotonin reuptake inhibitors. Int. J. Antimicrob. Agents 14, 177–180 (2000).

60. Bohnert, J. A., Szymaniak-Vits, M., Schuster, S. & Kern, W. V. Efflux inhibition by selective serotonin reuptake inhibitors in *Escherichia coli*. J. Antimicrob. Chemother. 66, 2057–2060 (2011).

61. Li, L., Kromann, S., Olsen, J. E., Svenningsen, S. W. & Olsen, R. H. Insight into synergetic mechanisms of tetracycline and the selective serotonin reuptake inhibitor, sertraline, in a tetracycline-resistant strain of *Escherichia coli*. J. Antibiot. (Tokyo). 70, 944–953 (2017).

62. Peterson, E. J. R., Ma, S., Sherman, D. R. & Baliga, N. S. Network analysis identifies Rv0324 and Rv0880 as regulators of bedaquiline tolerance in *Mycobacterium tuberculosis*. Nat. Microbiol. 1, 16078 (2016).

63. Brochado, A. R. et al. Species-specific activity of antibacterial drug combinations. Nature 559, 259–263 (2018).

64. Worthington, R. J. & Melander, C. Combination approaches to combat multidrug-resistant bacteria. Trends Biotechnol. 31, 177–184 (2013).

65. Weinstein, Z. B., Bender, A. & Cokol, M. Prediction of synergistic drug combinations. Curr. Opin. Syst. Biol. 4, 24–28 (2017).

66. Levin-Reisman, I. et al. Antibiotic tolerance facilitates the evolution of resistance. Science (80-.). 355, 826–830 (2017).

67. Loui, C., Chang, A. C. & Lu, S. Role of the ArcAB two-component system in the resistance of *Escherichia coli* to reactive oxygen stress. BMC Microbiol. 9, 183 (2009).

68. Cokol-Cakmak, M., Bakan, F., Cetiner, S. & Cokol, M. Diagonal method to measure synergy among any number of drugs. JoVE (Journal Vis. Exp. e57713 (2018).

69. Baldi, P. & Long, A. D. A Bayesian framework for the analysis of microarray expression data: regularized t-test and statistical inferences of gene changes. Bioinformatics 17, 509–19 (2001).

70. Huang, D. W., Sherman, B. T. & Lempicki, R. A. Systematic and integrative analysis of large gene lists using DAVID bioinformatics resources. Nat. Protoc. 4, 44 (2008).

71. Peterson, E. J. R. et al. Path-seq identifies an essential mycolate remodeling program for mycobacterial host adaptation. Mol. Syst. Biol. 15, (2019).

72. Arrieta-Ortiz, M. L. et al. An experimentally supported model of the *Bacillus subtilis* global transcriptional regulatory network. Mol. Syst. Biol. 11, (2015).

73. Keseler, I. M. et al. The EcoCyc database: reflecting new knowledge about *Escherichia coli* K-12. Nucleic Acids Res. 45, D543–D550 (2017).

74. Orth, J. D. et al. A comprehensive genome-scale reconstruction of *Escherichia coli* metabolism—2011. Mol. Syst. Biol. 7, (2011).

